# Ventromedial striatal dopamine dynamically integrates motivated action and reward proximity

**DOI:** 10.64898/2025.12.02.691784

**Authors:** Eugenia Z. Poh, Nicky L. Buitelaar, Gino Hulshof, Lucie Mazé, Pieter N. de Greef, Georgios Zaverdinos, Ingo Willuhn

## Abstract

Dopamine release in the ventromedial striatum (VMS) both invigorates actions and encodes reward-related information, yet how these functions are integrated remains under active debate. To investigate this further, we designed four different versions of a rat Go/No-go task, where we systematically manipulated response requirements, temporal task demands, and controllability of reward pursuit. Dopamine release increased reliably during action initiation (Go) but was delayed during action suppression (No-go), and this difference was insensitive to augmented response demands or controllability. Following response completion, dopamine rose gradually until animals arrived at the reward location, irrespective of reward-delivery timing, prior action demands, or controllability. This proximity dopamine-signal was exaggerated after animals exhibited Pavlovian consummatory behavior during No-go trials, revealing a motivational signal component. Together, these findings indicate that in reward contexts, VMS-dopamine signals successively integrate the invigoration of action initiation with the continuous tracking of spatial - but not temporal - proximity to rewards.

## Introduction

Appropriate, situation-dependent behavior in everyday life requires both motor control and information about upcoming rewards. A key neuromodulator involved in these processes is dopamine. Midbrain dopaminergic neurons have been shown to encode a reward-prediction error (RPE) which signals the discrepancy between expected and obtained outcomes in a given situation (Bayer and Glimcher, 2005; Montague et al., 1996; Schultz et al., 1997; Starkweather and Uchida, 2021; Steinberg et al., 2013). This reward-learning signal can also be observed in dopamine release from terminals in the ventromedial striatum (VMS) (Day et al., 2007; Kim et al., 2020; Mikhael et al., 2022; van Elzelingen et al., 2022), one of the main projection targets of midbrain dopamine neurons. Beyond reward learning, VMS dopamine is also implicated in invigorating actions toward appetitive rewards (Coddington and Dudman, 2018; Collins et al., 2016; Hamid et al., 2016; Howe et al., 2013; Howe and Dombeck, 2016; Mohebi et al., 2019; Nicola, 2007; Phillips et al., 2003; Roitman et al., 2004; Syed et al., 2016), supporting the view that the VMS acts as a motivation-action interface (Mogenson et al., 1980; Salamone and Correa, 2012, 2002). For example, VMS dopamine release increases upon movement towards a known reward location, or when initiating an instrumental response for rewards (Hamid et al., 2016; London et al., 2018; Mohebi et al., 2019; Phillips et al., 2003; Roitman et al., 2004; Syed et al., 2016). Although reward learning and motivation have been widely studied independently , how these two functions are integrated in the VMS remains unclear.

An additional process that influences striatal dopamine is the execution of actions required for achieving a reward. For example, under some conditions, the action-related *effort* required for a reward alters VMS dopamine (Cousins et al., 1996; Gan et al., 2010; Salamone and Correa, 2024). Furthermore, VMS dopamine responds differentially to reward contingencies across instrumental- and Pavlovian-conditioning tasks (Goedhoop et al., 2023; Hamid et al., 2021), and midbrain single-neuron activity differs reportedly for actions that are triggered externally (cue-initiated) versus ones that are under exclusive self control (self-initiated) (Romo and Schultz, 1990). Thus, these data suggest that reward-related VMS dopamine signals may be modulated by the effort or the controllability of reward seeking.

Further evidence for a role of VMS dopamine in motivation comes from the gradual increase in its release as animals approach a known reward location (Hamid et al., 2016; Howe et al., 2013; Kim et al., 2020; Mohebi et al., 2019). This signal scales with spatial proximity to rewards (Howe et al., 2013; Kim et al., 2020), and this prolonged “ramp”-like signal has been suggested to sustain motivational drive (Howe et al., 2013), reinforcing the preceding actions that led to reward (Hamid et al., 2016; Niv and Schoenbaum, 2008). However, such evidence is still limited, and it is unknown whether this gradual increase as animals approach the reward location is modulated by effort and the controllability of reward pursuit, and whether the signal is affected by the preceding action requirement for earning rewards.

In both human and rodent studies, Go/No-go tasks are commonly employed to study the effects of action initiation and suppression. However, action requirements for ‘Go’ and ‘No-go’ trials vary substantially between studies. Here, we developed multiple Go/No-go task variants to elucidate how VMS dopamine integrates reward-related information, action selection, effort and controllability of reward pursuit, and reward approach. More specifically, rats were trained to initiate (Go) or suppress (No-go) actions for rewards in response to an action-cue. In another task variant, we added ‘Free’ trials that required no overt action, which is akin to human ‘No-go’ trials where participants must withhold action for a reward (e.g., not press a button). To determine whether the dopamine differential between action and action suppression is influenced by reward-related contingencies, we manipulated: 1) controllability of reward seeking, defined as the ability to determine when to initiate reward pursuit (Self- vs Cue-initiated trials), 2) trial demands, and 3) timing of reward delivery. We hypothesized that controllability and effort would modulate the dopamine differential between action and action suppression, while the gradual increase in VMS dopamine during reward approach would generalize across contingencies.

By combining real-time measurements of dopamine release with detailed behavioral analysis, we replicate previous findings demonstrating that movement is critical for reward-related dopamine release (Syed et al., 2016). In addition, we demonstrate that VMS dopamine is governed specifically by the initiation of the approach to the reward location and proximity to this location, irrespective of preceding action requirements. Finally, different behavioral strategies of action suppression modulate dopamine release amplitude related to spatial reward proximity. Consummatory Pavlovian behaviors during action suppression are associated with the largest dopamine responses during subsequent reward approach, suggesting that individual differences in Pavlovian drive modulate dopamine encoding of spatial proximity to reward.

## Results

To investigate the role of VMS dopamine in reward processing and motor control, we implanted rats with fast-scan cyclic voltammetry (FSCV) microelectrodes (*n* = 21; Figure 2a) and trained them in four variants of the task: 1) a “short-task variant”, in which rats *Self-initiated* trials, 2) a “short-task variant” in which trials were *Cue-initiated* (altered controllability of reward pursuit), 3) a “long-task variant” of the *Self-initiated* task, in which the required duration of rat response action was increased (altered difficulty), and 4) a “long-task variant” of the *Cue-initiated* task (Figure 1a, see *Methods: Behavioral procedures*). We defined controllability as the rats’ ability to choose the time point of reward pursuit. In *Cue-initiated* trials, the time point at which trials could be started was dictated by a cue light, whereas in *Self-initiated* trials rats were able to choose intrinsically (control) when to attempt a trial start. There were two trial types in the short-task variants of the task: depending on the (auditory) action cue, rats were required to press a lever (Go) or nose poke (No-go) to obtain reward. In the short-task variants, action-cues switched off after trial completion and a reward was delivered immediately. In the long-task variants, we introduced a third trial type: in addition to Go and No-go action-cues and corresponding required responses, a third (auditory) cue signaled that reward would be delivered without requiring an overt action (Free trials). In the long-task variants, action-cues switched off after trial completion, or in the case of Free trials after 3s, and reward was delivered 2s later (Figure 1a).

**Figure 1.**
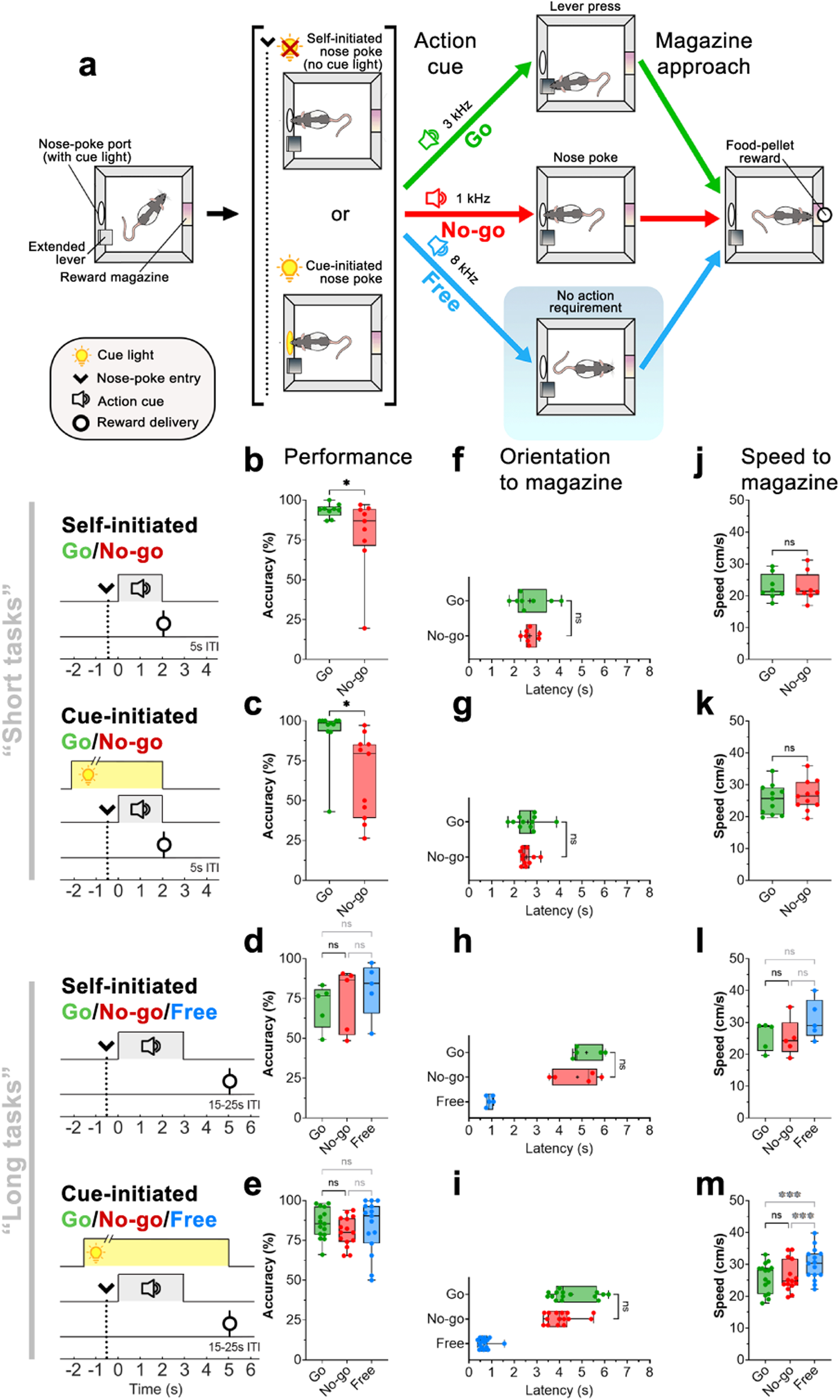
**Go and No-go performance is similar across task variants. *a)*** *Top:* Schematic of the Go/No-go (short) behavioral task. Rats initiated trials by entering a nose-poke port (arrow, dotted line), either voluntarily (*Self-initiated)* or following nose-poke light illumination (*Cue-initiated*). After staying in the nose-poke port for 0.5s, an auditory stimulus (action cue) instructed rats to either initiate action (lever presses: ‘*Go*’, green) or remain in the nose-poke port (‘*No-go*’, red). In the *Go/No-go/Free* (long) task, an additional action cue was presented that did not require an action for rewards (‘*Free*’, blue). Following Free cue onset, rats could approach the reward magazine immediately. *Bottom:* Schematic of task timeline. For all trials, reward-delivery latency (relative to action-cue onset) was similar. Shaded gray area depicts approximate duration of action-cue onset for “short” and “long” task variants ***b-e)*** Success rates across Go/No-go task variants. ***f-i)*** Latency to orient toward the reward magazine following action-cue onset. ***j-m)*** Speed of movement toward the reward magazine after orientation. Significance: * p < 0.05, ** p < 0.01, *** p < 0.001, Wilcoxon matched-pairs signed rank or *post hoc* Dunn’s tests.

**Figure 2.**
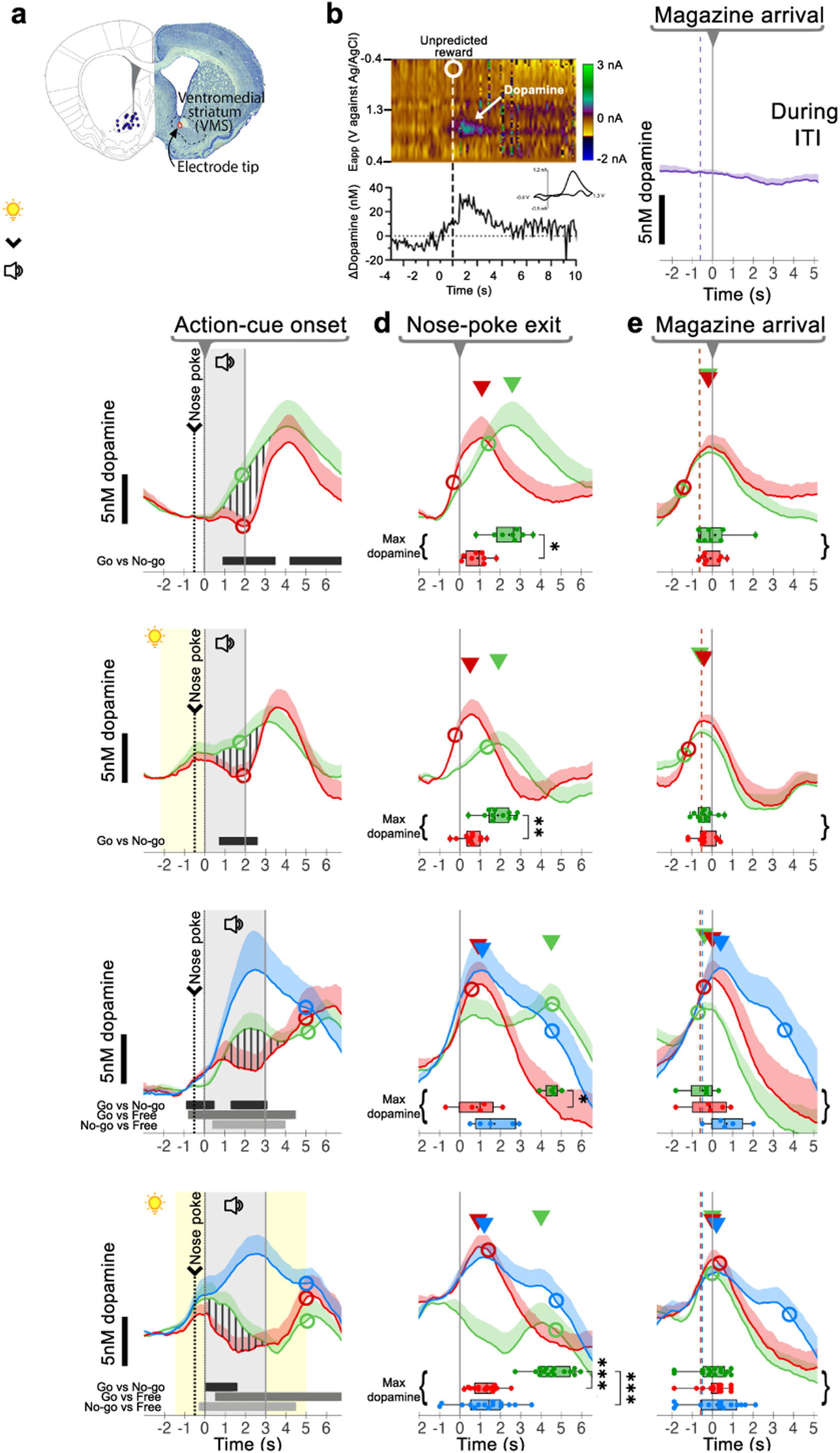
**VMS dopamine encodes both action initiation and reward-magazine arrival irrespective of action contingency. *a)*** Histological verification summary of VMS electrode placements for all animals used in this study (n = 21). **b)** Representative color plot and corresponding dopamine trace in response to an unpredicted pellet. The circle indicates pellet delivery. Inset: Accompanying cyclic voltammogram confirms dopamine detection. ***c-e)*** Average dopamine concentration (nM; mean + SEM) for Go (green), No-go (red), and Free (blue) trials. Triangles denote the average latency-to-maximum dopamine relative to different behavioral epochs. Subject-wise comparisons of dopamine data were made for all alignments. ***c)*** Traces aligned to action-cue onset. Shaded gray area depicts approximate duration of action-cue onset for “short” and “long” task variants. Vertical stripes highlight differences in Go vs. No-go dopamine. Horizontal bars indicate timepoints with significant within-subjects differences between trial types . ***d)*** Traces aligned to nose-poke exit and ***e)*** magazine arrival were analyzed by comparing latency-to-max dopamine release. Box plots depict individual latencies to maximum dopamine release (± 2s) around magazine arrival for each animal. ***f)*** Dopamine release during the inter-trial interval (ITI) as rats approached the reward magazine from the opposite wall, aligned to magazine arrival. Boxplot statistical significance: * p < 0.05, ** p < 0.01, *** p < 0.001, Wilcoxon matched-pairs signed rank or *post hoc* Dunn’s tests.

### Behavioral performance in Go/No-go (“short task”) was unaffected by controllability of reward pursuit

In the *Self-initiated* Go/No-go task, similarly used previously by Syed et al. 2016, trials were initiated by an (uncued) nose poke (Figure 1a). Our task design ensured that the time of reward delivery following action-cue onset was matched across trial types. After animals achieved proficiency in the task (see Methods), we measured behavioral performance and real-time dopamine release. Animals could initiate a trial by entering the nose-poke port after the intertrial interval (ITI); importantly, rats spent <20% of the ITI time inside the nose-poke port (Supplementary Figure S1), thus in between trials they mostly left the port. Reward success rates were significantly higher when animals had to initiate actions (Go trials) as compared to trials where the animals had to suppress actions (No-go trials) (Figure 1b; Wilcoxon matched-pairs test, W = 35, p = 0.039).

The activity of midbrain dopaminergic neuronal activity differs depending on the controllability of reward pursuit (Romo and Schultz, 1990). To evaluate how VMS dopamine dynamics during action initiation and suppression are affected by diminished control over trial initiation, we additionally trained our rats on a *Cue-initiated* task variant in which trials could only be initiated (by nose poke) after a cue turned on (Fig 1a;): rats learned to promptly respond to the cue (nose-poke light illumination) to initiate trials (1.68 ± 0.28 s; Supplementary Figure S2), indicating that they learned the contingency. Similar to the *Self-initiated* variant, rats trained in the *Cue-initiated* task showed significantly higher success rates on Go trials as compared to No-go trials (Figure 1c, *Cue-initiated* Go/No-go; W = 54, p = 0.014).

### Behavioral performance in Go/No-go/Free (“long task”) was unaffected by controllability of reward pursuit

Previous research has shown that spatial proximity of subjects relative to the reward location is reflected in VMS dopamine concentration (Howe et al., 2013), and dopamine responses may differ depending on whether rewards are contingent on a subject’s behavior (i.e., Pavlovian vs. instrumental responding for rewards) (Goedhoop et al., 2023; Hamid et al., 2021). Furthermore, VMS dopamine may be influenced by effort (Cousins et al., 1996; Gan et al., 2010; for review, see Salamone and Correa, 2024). To investigate the influence of these factors, we trained rats on the long-task variants that included ‘Free’ trials and increased Go and No-go trial requirements (Go: “long” repeated presses, 6.52 ± 0.32 presses, vs. “short” presses, 1.80 ± 0.03s; No-go: “long” vs. “short” nose-poke hold; see *Methods*, Figure 1a). Reward-delivery timing in Free trials matched the timing of long-task Go and No-go trials. In this *Cue-initiated* variant, rats promptly responded to the cue (0.95 ± 0.04 s, Supplementary Figure S1), indicating that they understood the contingency. Success rates did not differ significantly across Go/No-go/Free trial types for both *Self-initiated* (Figure 1d; Kruskal-Wallis test H(2) = 0.400, p = 0.954) and *Cue-initiated* (Figure 1e, H(2) = 1.458, p = 0.482) task variants.

### Motivation to approach the reward magazine was similar between Go and No-go trials

To determine whether action initiation (Go) and suppression (No-go) affected motivational drive to approach the reward location, we tracked behavioral performance using DeepLabCut (Mathis et al., 2018). In the short *Self-* and *Cue-initiated* Go/No-go task variants, the latency to orient to the reward magazine (Figure 1f; *Self-initiated*: W = 0, p > 0.999; Figure 1g; *Cue-initiated*: W = 6, p = 0.831), approach speed to the magazine (Figure 1j; *Self-initiated*: W = −5, p = 0.773, Figure 1k; *Cue-initiated*: W = −28, p = 0.240), and latency to arrive at the magazine after action-cue onset (Supplementary Figure S3; *Self-initiated*: W = −4, p = 0.844; *Cue-initiated*: W = 2, p = 0.966) did not differ between Go/No-go trial types.

To assess whether 1) altered controllability of reward pursuit, 2) increased task effort, and 3) the addition of Free trials affected the *core* comparison between action initiation (Go) and suppression (No-go), we examined behavioral performance in the Go/No-go/Free task variants. Trial type significantly influenced latency to orient to the reward magazine in both Go/No-go/Free task variants (Figure 1h; *Self-initiated*: H(2) = 7.600, p = 0.024; Figure 1i; *Cue-initiated*: H(2) = 24.13, p < 0.001). *Post hoc* Dunn’s test revealed orienting latencies did not significantly differ between Go and No-go trials (all p > 0.05, Figure 1h, k), but that Free trials significantly differed from Go and No-go trials (*Self-initiated* Go vs. Free: p = 0.034; *Cue-initiated*: p < 0.001, No-go vs. Free: p = 0.002), which was consistent with Free trials requiring no action compared to Go (action) and No-go trials (action suppression). Speed to approach the reward magazine was not significantly different for the *Self-initiated* task (Figure 1l; H(2) = 2.800, p = 0.367) but differed for the *Cue-initiated* Go/No-go/Free task (Figure 1m; H(2) = 19.73, p < 0.001). *Post hoc* Dunn’s test revealed that rats moved at similar speeds between Go and No-go trials (*Cue-initiated*), but were significantly faster on Free trials compared to Go and No-go trials (see Figure 1m; p < 0.01), suggestive of increased vigor in Free trials despite matching reward delivery times. In addition, although the latency to arrive at the reward magazine was significantly different between trial types for *Self-* (H(2) = 7.600, p = 0.024) and *Cue-initiated* Go/No-go/Free *tasks* (H(2) = 22.80, p < 0.001), *post hoc* Dunn’s test revealed no difference in arrival latencies between Go and No-go trials, but did earlier arrival for Free trials (Supplementary Figure S3). Together, these results indicated that Free trials were associated with earlier approach to the reward location, but the motivational drive to approach was comparable between Go and No-go trials, and was not affected by reward controllability (self- and cue-initiated) nor increasing task effort (short and long-task variant).

### VMS dopamine release encodes reward-related action initiation

VMS dopamine recordings (aligned to action-cue onset; Figure 2a,b) showed a rapid increase following presentation of the Go-action cue but no equivalent increase in response to the No-go-action cue (Figure 2c): Within-subject time-series comparisons (Jean-Richard-dit-Bressel et al., 2022) of Go and No-go trials (Figure 2c, ‘*Self-initiated* Go/No-go’, vertically striped area) confirmed significant differences (*p* < 0.05; Figure 2c, thick horizontal bar insets). Despite differences in task requirements such as controllability to attain reward (i.e., *Self-* vs. *Cue-initiated*) and increased effort for Go and No-go trials in the Go/No-go/Free task (i.e., long-task variant, see *Methods*, Figure 1a), within-subject comparisons for all task variants showed a consistent, significant increase in dopamine release on Go trials compared to No-go trials (*p* < 0.05; Figure 2c *Self-* and *Cue-initiated* Go/No-go/Free task, thick horizontal bar insets). In addition, dopamine release was highest for Free trials as compared to Go and No-go trials (p < 0.05; Figure 2c thin horizontal bar insets). Our data suggested that the increase in VMS dopamine release not only reflected reward-predictive cues but also action initiation. In addition, the difference in VMS dopamine during action initiation (Go) and suppression (No-go) was unaffected by controllability of reward pursuit and increased trial demands.

### VMS dopamine release does not only reflect reward-related action initiation

Since the rise in dopamine release for Go trials occurs before rises in No-go trials when traces were aligned to action-cue onset (Figure 2c, vertical line at t=0s), we sought to clarify whether this difference may be due to an earlier moment of action initiation (nose-poke exit) for Go trials as compared to No-go trials. Therefore, we aligned traces to the time when animals departed the nose-poke port (Figure 2d). For Go trials, this was the moment when animals departed the nose-poke port to move to the lever, and for No-go trials, this was the moment animals successfully completed the trial. Analysis of maximum dopamine release (Figure 2d, triangles) differed between trial types: in contrast to No-go and Free trials, maximum dopamine release during Go did not align with nose-poke port exit (Figure 2d). In particular, the latency to maximum dopamine was significantly different between trials, where maximum dopamine release was consistently delayed for Go trials (Figure 2d, boxplot insets; *Self-initiated* Go/No-go: W = 34, p = 0.016; *Cue-initiated* Go/No-go: W = 64, p = 0.002; *Self-initiated* Go/No-go/Free: H(2) = 8.40, p = 0.008; *Cue-initiated* Go/No-go/Free: H(2) = 23.33, p < 0.001). Overall, our data suggested that differences in Go vs. No-go dopamine could not be attributed to the moment of action initiation (nose-poke departure) alone.

### Maximum VMS dopamine release encodes spatial but not temporal proximity to rewards

Previous research has shown that VMS dopamine can not only encode RPE and movement, but can also reflect a subject’s proximity to the expected reward location (Hamid et al., 2016; Howe et al., 2013; Mohebi et al., 2019). We therefore realigned dopamine traces to the moment animals arrived at the reward magazine (Figure 2e, vertical line at t=0s). Across all trial types and sessions, dopamine release consistently peaked around the time of magazine arrival (Figure 2e, boxplot insets; *Self-initiated* Go/No-go: W = 7, p = 0.672; *Cue-initiated* Go/No-go: W = 3, p = 0.916; *Self-initiated* Go/No-go/Free: H(2) = 4.526, p = 0.111; *Cue-initiated* Go/No-go/Free: H(2) = 4.204, p < 0.122). When plotting maximum dopamine as a function of distance to the magazine, the distribution revealed a prominent peak around the magazine-panel (Supplementary Figure S10). Importantly, the timing of maximum dopamine (Figure 2e, *Self-* and *Cue-initiated* Go/No-go/Free, triangles) did not correlate with the timing of reward delivery: in Free trials, average reward-delivery latency (Figure 2e, hollow circles) was delayed following magazine arrival by up to 3s. Realignment of maximum-dopamine concentrations around the time of reward delivery showed that there was no peak distribution around this epoch (Supplementary Figure S11, in particular, see Free trials). In addition, rats oriented to the magazine at similar times, across trial-types and task variants, relative to magazine arrival (Figure 2e; on average 0.6s from orientation to magazine arrival; vertical dashed line). To determine whether maximum dopamine reflected the approach to the reward magazine, or required information about an upcoming reward (e.g., action cues), we analyzed changes in dopamine release during *unprompted* magazine entries in the *Self-initiated* Go/No-go/Free task (Figure 2f, *During ITI*). Our findings demonstrated that unprompted magazine entries during the ITI did not elicit changes in VMS dopamine release. Finally, to determine whether maximum dopamine primarily reflects the earliest predictor of reward, we realigned traces to the moment of action-cue onset in *Self-initiated* trials, and to nose-poke light illumination for *Cue-initiated* trials (Supplementary Figure S12). Our results showed that there was no consistent peak in the distribution of maximum dopamine around cue onset. Together, our results suggested that increases in VMS dopamine required task-relevant features such as knowledge of an upcoming reward and movement towards the reward magazine, where VMS dopamine was maximum upon arrival at the expected reward location.

### Motivational state reflected by No-go behavioral strategy correlates with dopamine signal size during reward approach

During the action-cue period of No-go trials, rats had to suppress actions by remaining in the nose-poke port. However, we observed individual differences in behavioral strategies in performing ‘action suppression’. We therefore classified No-go trials into three behavioral strategies (*Biting* (purple), *Digging* (teal), *Calm* (maroon); Figure 3a) using a decision-tree classifier based on No-go behavior (Supplementary Figure S4; Video 1-3). Dopamine concentrations during the action-cue period revealed no differences between classifications (Figure 3b, Supplementary Figure S5). This result indicates that behavioral strategy during No-go trials was unrelated to dopamine release *during* the action-cue period, as they demonstrated similar dopamine release for ‘action suppression’ despite varying No-go behavioral strategies.

**Figure 3.**
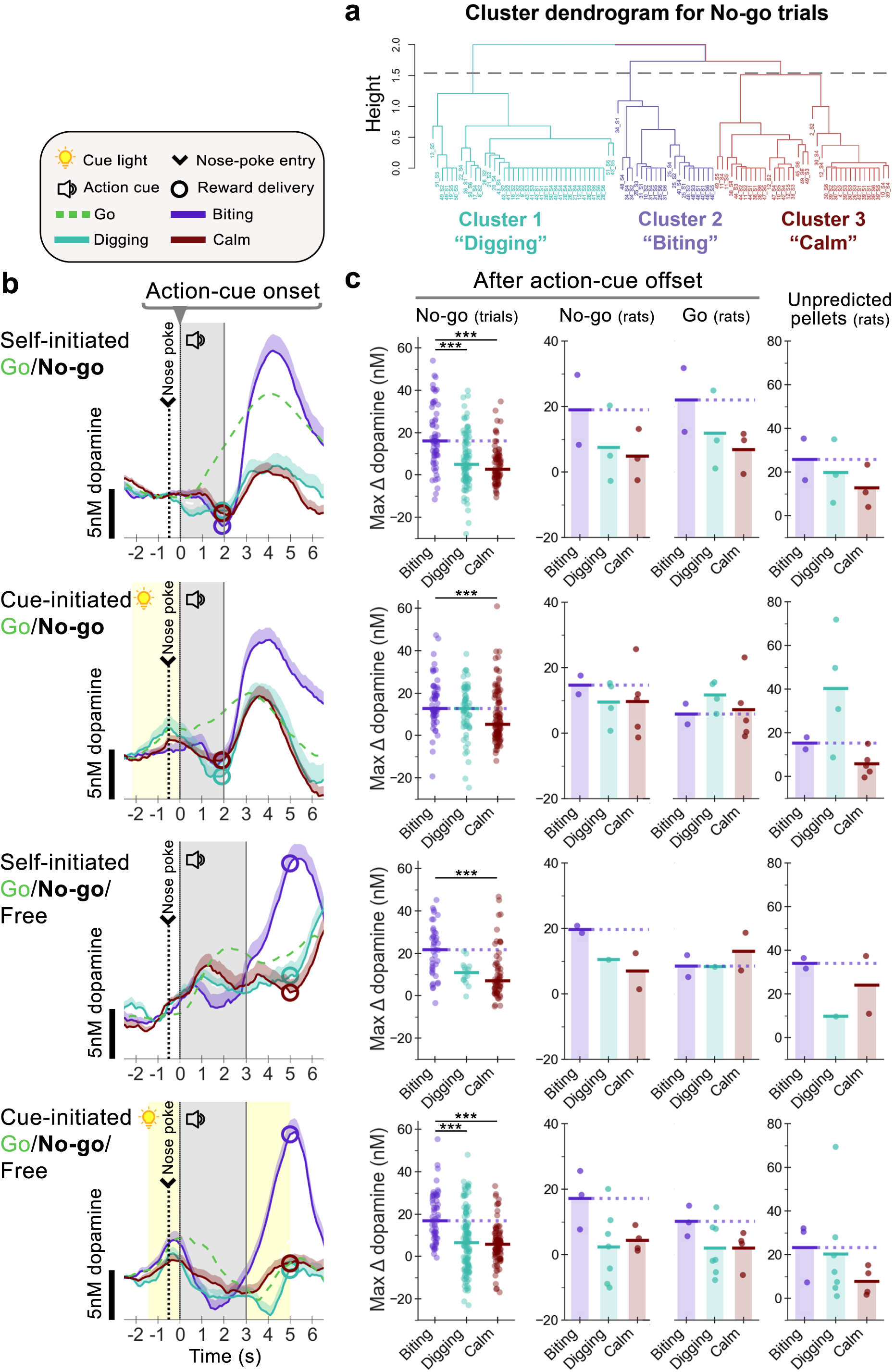
**Classified No-go behavior reveals VMS dopamine release following action-cue offset in *‘Biting’* trials. *a)*** *Left*: Agglomerative hierarchical clustering of behavior during the No-go action-cue period using Ward’s linkage and Euclidean distance. A three-cluster solution (dotted grey line) was selected resulting in a balanced distribution of trials across clusters and sessions. These clusters corresponded to three distinct behavioral strategies. *‘Biting’* - defined as biting of the nose-poke-port wall; *‘Digging’* - digging movements in the nose-poke port; *‘Calm’* - all other no-go trials. ***b)*** A supervised decision-tree classifier labeled trials to one of three behavioral groups: *Digging*, *Biting,* and *Calm*. Average dopamine concentration of each classified group is depicted (nM; mean + SEM). ***c)*** Maximum dopamine concentration (nM) following action-cue offset (+2s) was highest in *Biting* trials as compared to other No-go trials. *‘After action-cue offset: No-go (trials)’*: Individual No-go trials classified by No-go behavior, and trial-wise statistical Kruskal-Wallis tests were performed for each task-variant. Solid lines depict the median. *‘After action-cue offset: No-go (rats)’, ‘After action-cue offset: Go (rats)’,* and ‘*Unpredicted pellets (rats)*’: Rats classified based on their predominant No-go strategy, with no statistical tests performed. Solid lines depict the mean. Significance: * p < 0.05, ** p < 0.01, *** p < 0.001, *post hoc* Dunn’s tests.

However, maximum dopamine concentration measured *after* action-cue offset (Figure 3c) significantly differed between strategies for No-go trials in all task variants (*Self-initiated* Go/No-go: H(2) = 34.15, p < 0.001; *Cue-initiated* Go/No-go: H(2) = 16.24, p < 0.001; *Self-initiated* Go/No-go/Free: H(2) = 22.54, p < 0.001; *Cue-initiated* Go/No-go/Free: H(2) = 37.48, p < 0.001). *Post hoc* comparisons showed that dopamine levels were consistently higher for *Biting* as compared to *Calm* trials (Figure 3c, *‘After action-cue offset: No-go trials’*, all *p* < 0.001). Dopamine was also higher for *Biting* as compared to *Digging* in two task types (Figure 3c). Behaviorally, we observed that Biting trials were characterized by earlier nose-poke hold exits (Supplementary Figure S6). These findings suggest that while dopamine release during the action-cue period is unaffected by action-suppression strategy, differences emerged during reward approach and may reflect differences in motivational drive.

Regrouping animals based on their predominant No-go behavioral strategy revealed a trend in maximum dopamine that resembled individual trial data (Figure 3c, ‘*No-go (rats)*‘ and ‘*No-go (trials)*‘, respectively). Therefore, we grouped animals based on their predominant action-suppression strategy to determine whether *Biting* animals also showed elevated dopamine in response to other reward-related information. However, maximum dopamine release in response to an unpredicted pellet did not follow this pattern and instead, they were similar across all groups (Figure 3c, ‘*Unpredicted pellets (rats)*’). Comparably, grouped rats did not show differences in maximum dopamine for Go and Free trials following action-cue onset (Supplementary Figure S5). These results suggested that the elevated dopamine response during reward approach in *Biting* animals was specific to No-go trials.

To further explore whether VMS dopamine reflects motivational drive within another trial type, we examined Go trials split by the timing of the last lever-press (Supplementary Figure S7). Trials with later last lever-presses (slowest 25% of trials) showed lower maximum dopamine, an effect most pronounced in the long-task variant where lever presses could extend beyond the action-cue period. Together with our classified No-go trial analysis, these results suggested that behavioral indicators of motivational engagement (No-go: *Biting* and nose-poke hold latency; Go: last lever-press latency) modulated maximum dopamine during reward approach.

## Discussion

We examined how VMS dopamine integrates motivation and reward-related information during action initiation and suppression in four Go/No-go task variants. In our task, *action initiation* occurred at two distinct points after trial start: 1) when rats began lever pressing (Go), and 2) when rats walked to the reward magazine, either without action requirement (Free) or after successful trial completion (Go and No-go). By contrast, *action suppression* was unique to No-go trials, where animals were required to stay in the nose-poke port. Dopamine release during action (Go) was consistently higher than during action suppression (No-go); this difference was unaffected by controllability of reward pursuit (*Self-* vs. *Cue-initiated*) and differences in effort (i.e., short vs. long). Additionally, our data suggest that VMS dopamine signaled the subjective spatial proximity to reward location. This conclusion is supported by the fact that this gradual increase in dopamine persisted despite differences in the timing of reward delivery (i.e., with or without a 2s-delay after action), changes in controllability, and preceding action requirement. Our results indicate that in a reward context, VMS dopamine is not only associated with action execution but continuously tracks spatial, but not temporal, proximity to rewards. Unexpectedly, consummatory Pavlovian behavior in the form of (wall) biting during action suppression led to markedly larger dopamine release during subsequent reward-magazine approach than other behaviors. Together, our findings indicate that VMS dopamine sequentially integrates reward-driven action initiation with the gradual increase in spatial reward proximity.

### Reward-related dopamine depends on action initiation irrespective of controllability and effort

VMS dopamine has been associated with the invigoration and initiation of reward-seeking behavior (Ikemoto and Panksepp, 1999; Nicola, 2007; Robinson and Berridge, 1993) as demonstrated by both bulk measurements of VMS dopamine release (Collins et al., 2016; du Hoffmann and Nicola, 2014; Hamid et al., 2016; Mohebi et al., 2019; Nicola, 2010; Phillips et al., 2003; Roitman et al., 2004; Syed et al., 2016; Wassum et al., 2012) and the activity patterns of individual midbrain dopamine neurons (Coddington and Dudman, 2018; Jin and Costa, 2010). Across all four of our task variants, VMS dopamine release was consistently higher during action (Go) than during action suppression (No-go). Thus, by extending the pioneering work by Syed and colleagues (Syed et al., 2016), our findings provide compelling evidence that reward-related dopamine release in the VMS is a function of associated reward-driven action.

In addition to action initiation, recent work suggests that striatal dopamine dynamics are shaped by the controllability of reward contingencies (Bernklau et al., 2024; Goedhoop et al., 2023; Hamid et al., 2021). For instance, the anticipation of operant action initiation during reward-cue presentation is accompanied by more sustained VMS dopamine release compared to cues that signal reward delivery without action requirement (Goedhoop et al., 2023). Similarly, midbrain dopamine recordings showed increased responses to externally triggered actions (i.e., *Cue-initiated*) than to *Self-initiated* actions for rewards (Romo and Schultz, 1990). In contrast, our data indicate that manipulations of controllability in reward pursuit (i.e., *Self-* vs. *Cue-initiated* trials) did not affect the contrast between Go and No-go dopamine dynamics during action execution, but instead only elicited dopamine release at an earlier time point in the trial, presumably triggering an earlier positive RPE in *Cue-initiated* trials (Figure 2a). This discrepancy to previous studies may be rooted in differences in experimental design. Goedhoop and colleagues (2023) compared VMS dopamine during operant versus Pavlovian conditioning, similar to the comparison of Go and Free trials in our task, respectively, which also display differences in VMS dopamine (Figure 2c). On the other hand, the discrepancy may also be rooted in different definitions of “controllability”. Here, we defined controllability as the moment animals can initiate reward pursuit (i.e., self- vs cue-initiated), which considers a different aspect of controllability. Thus, we cannot exclude the possibility that dopamine would respond to a more extreme manipulation of controllability Indeed, Bernklau and colleagues (2024) showed that striatal dopamine reflects the animal’s perceived locus of control, which shifts from external cues toward the animal as the causal agent with experience. In our paradigm, both self- and cue-initiated task variants may have been equally perceived as agent-controlled (albeit curtailing timing and source of action initiation), but potentially leaving the inferred locus of control unchanged. Together, our results indicate a prominent role for VMS dopamine in reward-related action (Coddington and Dudman, 2018; Ikemoto and Panksepp, 1999; Nicola, 2007) that is not sensitive to moderate changes in task controllability.

Previous studies have shown that dopamine signals may be influenced by the effort required to obtain reward but only for particular task conditions (Cousins et al., 1996; Gan et al., 2010; see for reviews, Salamone and Correa, 2024; Walton and Bouret, 2019). We tested this by varying response demands: short and long Go trials requiring either two lever presses or repeated pressing for three seconds, respectively. No-go trials required matching two or three seconds of action suppression, respectively (Fig 1b). However, such effort manipulations had no detectable impact on action-related dopamine: the relative contrast in VMS dopamine in Go versus No-go trials did not differ between short and long task variants, suggesting that dopamine does not scale with response duration or physical effort. This is consistent with some previous work (Gan et al., 2010; Walton and Bouret, 2019).

The addition of Free trials in the current study provides critical insight into the role of dopamine in motivated action directed at reward. In Go/No-go tasks designed for human and non-human primates, Go trials often require an overt action (e.g., pressing a button) and No-go trials require passive inaction (e.g., not pressing the button; e.g., Raud et al. 2020). In the present study, Go and Free trials were akin to human Go and No-go trials, respectively. Importantly, our findings demonstrate that VMS dopamine dynamics are highly distinct between action initiation (Go), active inaction (No-go), and passive inaction (Free), for rewards. Therefore, precise definitions of “No-go” trials in both human and animal models are necessary for cross-species translatability of the neural basis of motivated behavior.

### VMS dopamine release reflects spatial proximity during approach of reward location

Beyond physical effort, the long-task variants imposed a two-second reward delay after action-cue offset, permitting clearer separation of dopamine signaling during action-cue and reward-approach epochs. Critically, maximum dopamine responses across all four task variants, could not be explained solely by action initiation, as the latency to maximum dopamine differed between trial types when aligned to nose-poke exit (Figure 2d, triangles), diverging from previous reports (Syed et al., 2016). Instead, dopamine release during the long-task variant revealed a gradual increase during magazine (reward location) approach that peaked upon arrival at the magazine, irrespective of whether rewards were received immediately or were delayed (Figure 2d). Importantly, we did not observe an increase in VMS dopamine when animals approached and checked the magazine between trials, indicating the necessity of reward expectation linked to trial performance (Figure 2e). In addition, the signal cannot be attributed to orientation to the reward-magazine - if orientation were the cause, dopamine would not be expected to increase while animals maintain a constant orientation when moving toward the reward magazine. Overall, this finding is consistent with several studies reporting a gradual rise in VMS dopamine, frequently referred to as ramps, as animals actively (Farrell et al., 2022; Hamid et al., 2016; Howe et al., 2013; Krausz et al., 2023; Mohebi et al., 2019) or passively (Farrell et al., 2022; Guru et al., 2020; Kim et al., 2020; Mikhael et al., 2022) move toward reward. This phenomenon was particularly pronounced during Free trials, where dopamine peaked when rats arrived at the reward magazine up to five seconds before reward delivery (Figure 2c,d Go/No-go/Free, circles). Together, our data suggest that while VMS dopamine initially facilitates the initiation of reward-driven action, it subsequently tracks progression toward a reward-associated location, which may serve to sustain motivational drive during reward approach.

The gradual increase in VMS dopamine during reward approach, peaking upon arrival, has been suggested to reflect ongoing state estimation under uncertainty (Farrell et al., 2022; Gershman and Uchida, 2019; Kim et al., 2020; Mikhael et al., 2022). Alternatively, this gradual increase, or ramp, has been suggested to reflect an ‘internal model’ that continuously updates reward-proximity estimates as animals progress toward the goal (Farrell et al., 2022; Guru et al., 2020). Our findings support the proximity account. In our Go/No-go task, rats traversed the same familiar, relatively short path to the magazine in thousands of trials, minimizing state uncertainty and reducing the need for ongoing state estimation updates. Instead, our findings suggest that VMS dopamine maintained a continuous estimate of spatial reward proximity: arrival at the reward magazine following trial completion served as the most reliable predictor of reward and coincided with maximum dopamine release, consistent with previous findings (Hamid et al., 2016; Howe et al., 2013; Mohebi et al., 2019). While our results clearly demonstrate dopamine signaling related to spatial proximity to rewards, our experimental design cannot definitively distinguish whether VMS dopamine encodes RPE computations or spatial proximity to sustain motivated approach behavior. More modeling and experimental work is needed to test these predictions (but see, Gershman et al., 2024; Lerner et al., 2021).

### Dopamine dynamics are linked to motivated action

Our findings reveal that VMS dopamine follows a consistent temporal pattern across trials: dopamine increases when animals initiate reward-directed actions, then continuously tracks spatial reward proximity. Critically, the magnitude of this proximity signal was modulated by motivational engagement, measured by trial-type-specific behavioral indicators (i.e., Go lever-press latencies and No-go behavioral strategies) and cannot be solely explained by changes in vigor. Go trials, in which lever-press requirements were completed faster, were associated with an earlier magazine approach and a larger and earlier increase in dopamine during the subsequent magazine approach than Go trials exhibiting slower action completion (Supplementary Figure 7). In No-go trials, we identified a distinct behavioral phenotype: consummatory Biting responses directed at the wall around the nose-poke device during action suppression were associated with shorter nose-poke hold latencies (Supplementary Figure S6) and an approximately two-fold increase in dopamine during subsequent reward approach, as compared to non-biting strategies across all task variants (Figure 3b). Notably, the increased dopamine signal (associated with earlier lever-press completion (Go) and Biting (No-go)) was specific to reward approach, and was not related to the action-cue period itself (Supplementary Figure S5-S7). Two lines of evidence argue against (Go and No-go) action vigor explaining these dopamine changes. First, the modulation of reward-approach dopamine occurred after the action-cue period, dissociating it temporally from any action vigor expressed during lever-pressing or nose-poke holding. Second, magazine approach-speed, which is a direct measure of post-action-cue vigor (i.e., behavior occurring at the same time as the observed differences in dopamine), showed no consistent relationship with dopamine magnitude across trial classifications (Supplementary Figures S5–S7).

Although Biting responses, which resembled consummatory behaviors typically directed toward rewards, shared similarities with so-called ‘sign-tracking’ behavior (Flagel et al., 2009; Hearst and Jenkins, 1974), several features distinguish our observations from sign tracking. First, sign tracking is promoted when reward-predictive cues and reward location are in spatial proximity (Christie, 1996; Flagel et al., 2009). However, in our experimental setup, reward-related manipulanda (i.e., levers, nose-poke port) and reward magazine, were positioned on opposing walls of an operant chamber exceeding standard size (Figure 1a). Second, it has been reported that rats do not readily sign-track to nose-poke ports or auditory cues (Beckmann and Chow, 2015; Meyer et al., 2014), both critical for No-go trials in our paradigm. Third, prior work demonstrated enhanced VMS dopamine during Pavlovian behavior towards the cue (Flagel et al., 2011, 2009), whereas we observed amplified dopamine *after* action suppression (and after the cue presentation), during reward approach.

These distinctions suggest that Biting may reflect elevated Pavlovian drive and/or difficulty suppressing reward-directed action, likely due to enhanced motivation (Robinson et al., 2014), rather than sign tracking. Accordingly, our findings point at inter-individual differences in behavioral strategy, analogous to sign and goal-tracking behaviors (Flagel et al., 2011; Robinson et al., 2014). Animals exhibiting heightened task engagement or Palovian drive showed amplified dopamine responses to identical changes in spatial reward proximity, suggesting that encoding of spatial-reward proximity is dynamically modulated by motivational state.

A potential explanation for differences in VMS dopamine may be attributed to Pavlovian biases, wherein reward-associated cues promote approach over suppression, typically resulting in higher Go than No-go performance accuracy. We indeed observed this behavioral asymmetry in our standard Go/No-go task; however, it disappeared when Free trials were included (Figure 1c). Importantly, VMS dopamine continued to differentiate between Go and No-go trials despite the absence of this phenomenon (i.e., more VMS dopamine during the action-cue period for Go than No-go; Figure 2c). Therefore, our data suggest that Pavlovian bias cannot account for the observed differences in dopamine dynamics.

### Conclusions

The findings of the present study are consistent with a general role of VMS dopamine in both RPEs and action. However, we further demonstrate that the differential between action and action suppression is not influenced by behavioral controllability or effort. Rather, our results suggest that VMS dopamine primarily encodes the initiation of reward-seeking actions and spatial proximity to rewards, thereby bridging two traditionally separate fields investigating the roles of dopamine in movement and in reward processing, respectively. Building on previous reports that individual dopaminergic neurons multiplex sensory, motor and cognitive information (Engelhard et al., 2019; Kremer et al., 2020), our bulk dopamine-terminal measurements suggest this multiplexing is extended to the broader population level. These findings suggest that population-level VMS dopamine signals serve as a unifying mechanism, conveying information about both motivated action and reward to postsynaptic VMS neurons. We propose that within this framework, VMS dopamine links motor and motivational processes to ensure that actions are maintained until reward receipt.

## Methods

### 1. Animals

At the start of the experiment, adult male Long-Evans rats (250-350g, Janvier Labs) were individually housed on a reversed 12 h light/dark cycle (lights off from 08:00 to 20:00) with controlled temperature and humidity. Water was available *ad libitum*. Following surgical procedures and one week of recovery, rats were food-restricted to 85% of their free-feeding body weight. All animal procedures were performed during the dark phase (between 11:00-19:00) in accordance with Dutch and European law, and approved by the Animal Experimentation Committee of the Royal Netherlands Academy of Arts and Sciences. A total of 21 rats were used in the experiments described in the following sections, where each rat had at least one functional and histologically verified recording electrode.

### 2. Stereotaxic surgery

Stereotaxic surgery was performed as previously described (Willuhn et al., 2012). Rats were given an analgesic (Metacam, 1-2 mg/kg, s.c.), anesthetized with isoflurane (2-3% maintenance), and placed in a stereotactic frame. Body temperature was maintained at 37°C during surgery using a heating pad and monitored with a probe. The scalp was shaved, disinfected with 70% alcohol, and a midline incision was made. The incision site was treated with lidocaine (100 mg/ml) and the periosteum gently removed to expose the cranium. Holes were drilled for 3-4 anchor surgical screws, two custom-made carbon-fiber microelectrodes(Clark et al., 2010) unilaterally (right hemisphere) targeting the VMS (A-P: +1.2, M-L: +1.5, D-V: −7.1 mm, relative to Bregma (Paxinos and Watson, 2006), and an Ag/AgCl reference electrode that was positioned separately in the forebrain. Electrodes were secured and anchored to the surgical screws with dental acrylic cement. Following surgery, rats were subcutaneously injected with 2 mL saline and placed in a temperature-controlled cabinet until they exhibited mobility. Following implantation of the electrodes, rats were individually housed and were left to recover for at least one week. The location of VMS electrodes can be found in Supplementary Figure S8.

### 3. Behavioral procedures

Three days before starting behavioral training in the operant boxes, 10-15 sugar pellets/day were introduced into the home cage. Behavioral experiments were performed in modified operant boxes (32 x 30 x 29 cm, MedAssociates Inc.), equipped with a nose-poke hole with an integrated cue-light, house light, auditory-tone generator, food-reward magazine, metal-check grid floor, and retractable levers (Figure 1a). Operant boxes were interfaced with a fast-scan cyclic voltammetry (FSCV) setup for dopamine recordings of awake and freely-moving animals and a video camera to survey behavior. The operant boxes were stored in individual Faraday cages insulated with sound-absorbing polyurethane foam and ventilated by a fan. Before commencing training or recordings in the operant boxes, rats were habituated to the room for at least 10 min.

#### Self-initiated task variant

##### Self-initiated Go/No-go task (“short”; n = 9)

Using a behavioral paradigm inspired by Syed and colleagues (Syed et al., 2016), rats initiated a trial by entering the nose-poke port for at least 0.5 s. One of two action (auditory) cues were played, instructing rats to either move towards and press on a unilaterally extended lever at least twice within 5 s (‘Go’; 3 kHz) or to remain in the nose-poke hole for another 1.7-1.9 s (‘No-go’; 1 kHz). After five correct Go trials, the contralateral lever extended and the opposing lever retracted. Successful trial completion was marked by the offset of the action cue and food pellet dispensed into the reward-magazine immediately. During training and recording sessions, trials were separated with a fixed intertrial interval (ITI) of 5 s.

##### Self-initiated Go/No-go/Free task (“long”; n = 5)

Go and No-go trial requirements were increased, together with the inclusion of a third trial type (long-task variant). During training, Go trials required an initial lever press within 1.7 s of action-cue onset and a subsequent lever press within the next 1.5 s. During recording days, a correct Go trial occurred when rats made the first lever press between 2.7 to 3.2 s after cue onset. For No-go trials, rats had to stay in the nose-poke hole for at least 2.7 to 3 s after cue onset. Following successful training of the extended Go and No-go trials, a “Free” trial was introduced (8 kHz tone). Rats were permitted to do anything but nose poke or lever press during the Free cue; most rats walked directly to the food-magazine following cue onset. Rewards were delivered with a delay of 2 s following successful trial completion. Trials were separated with a variable ITI between 15-25 s.

Failure to perform the trial’s required action resulted in house-light illumination for 5 s (i.e., incorrect trial). During training (i.e., non-recording days), the trial type was repeated to a maximum of three times or until the trial was correctly completed, whichever came first. During dopamine-recording days, trial type repetition did not occur. Success rates for each trial type was calculated by determining the percentage of correct/(correct+incorrect) trials, with incorrect trials only included if the error was made after action-cue onset.

Failure to hold the snout in the nose-poke port during the first 0.5 s before action-cue onset resulted in an immediate house-light illumination for 5 s and presentation of another ITI.

#### Cue-initiated task variant

The structure of *Cue-initiated* trials was similar to that of *Self-initiated* trials, except that they could only be initiated following the illumination of the nose-poke light. The nose-poke light stayed illuminated for up to 15 s until initiation, and remained illuminated until the end of the trial. All other trial requirements were identical, except that the house light illuminated for 5 s and a new ITI began when rats made a nose poke or lever press during the ITI, or if they did not perform a nose-poke within 15 s of nose-poke light illumination at the start of the trial. A total of *n* = 11 and *n* = 15 were included in the *Cue-initiated* Go/No-go and *Cue-initiated* Go/No-go/Free tasks, respectively.

### 4. Real-time dopamine recordings and analysis

Simultaneous dopamine and video recordings were carried out when rats were able to perform with >60% success rate for each trial type on at least two consecutive training sessions. On recording days, the trial types were counterbalanced. Within a session, Go left, Go right, and No-go trials were presented with 33% probability each, without replacement. This design results in overrepresentation of Go trials, which establishes the prepotent Go response. For sessions with Free trials, trials were presented with a 25% chance without replacement.

#### FSCV measurement and analysis

Fast-scan cyclic voltammetry (FSCV) at chronically implanted carbon-fiber microelectrodes (Clark et al., 2010) was used to record rapid changes in extracellular dopamine concentration. Prior to recording, microelectrodes were connected to a head-mounted voltammetric amplifier interfaced with a PC-driven data acquisition and analysis system (National Instruments) through an electrical commutator (Clark et al., 2010). Voltammetric scans were repeated every 100 ms (i.e., 10 Hz sampling rate). An alternating potential at the carbon-fiber electrode tip was applied from –0.4V versus Ag/AgCl to +1.3V (anodic sweep) and back to –0.4V (cathodic sweep) at 400V/s (total scan time of 8.5ms), and held at –0.4V between scans. Dopamine traces were isolated from the voltammetric signal by chemometric analysis using a standard training set, based on electrically stimulated dopamine release detected with chronically implanted electrodes (Clark et al., 2010). All data were smoothed with a 10-point median filter and baseline (set at 1s before event of interest, with the exception of 2.5 s before action-cue onset for *Cue-initiated* Go/No-go and Go/No-go/Free tasks) subtraction was performed on a trial-by-trial basis prior to analysis. Dopamine data were aligned to events of interest: action-cue onset, nose-poke exit, and magazine arrival.

#### Histological verification of recording sites

Following completion of behavioral and recording experiments, animals were terminally anesthetized using sodium pentobarbitone. An electrolytic lesion was performed by running a current on the microelectrodes to mark the FSCV recording sites. Rats were transcardially perfused with saline (0.9% NaCl, w/v) and paraformaldehyde (4% in PBS, w/v). Whole brains were collected and postfixed in Parafresh for 24 h, cryoprotected in sucrose solution (30% in PBS w/v), and cryosectioned coronally (40 μm). Slices were stained with cresyl violet. All animals included in this study had electrodes positioned in the ventromedial striatum (VMS; Supplementary Figure S8).

### 5. Video recordings and analysis

#### DeepLabCut pose estimation

For detailed quantification of behavior across each dataset, we employed DeepLabCut (DLC) to estimate body part coordinates of each animal (Mathis et al., 2018; see Supplementary Figure S9). A total of 1500 manually labelled frames from various videos were used to train a ResNet-50 based neural-network.

Labelled body parts included the snout, neck, four back-points, and tail-base. A reference frame from each video was labelled manually to determine the coordinates of the arena floor. Body coordinates were then transformed from pixels into cm using the ‘fitgeotform2d’ function in MATLAB. Custom analyses were developed in Python to filter DLC coordinates and to analyze behavioral features. Specifically, likelihood cut-off scores for each body part were set at 0.9, large movements in x and y coordinates were removed between each consecutive frame (> 1.5 cm between frames), body-part speeds exceeding a specific speed (mean speed of 9 consecutive frames + 2*SD), and linear interpolation was applied between the nearest valid x and y coordinates to fill in gaps in the data. Data then underwent Gaussian smoothing to reduce noise. This smoothing was performed using a Gaussian window with a SD calculated to capture approximately 99% of the Gaussian probability density function. The smoothing process was applied within a sliding window of 9 frames (29.97 fps) corresponding to approximately 300 milliseconds.

#### DeepLabCut behavior analysis

Orientation towards the magazine was determined by calculating the angle between the vector from the neck to the snout, and extending this vector to the wall containing the magazine-panel. Frames of animals were labelled as oriented when this vector crossed the length of the magazine-panel. Magazine arrival was determined as the moment the neck was <2 cm from the magazine-panel.

To analyze magazine approach during the ITI, we used DLC-derived coordinates to identify instances where animals were at least 25 cm away from the reward magazine and subsequently moved towards the magazine within 2s (similar to reward approach following a successful trial). This was performed in the *Self-initiated* Go/No-go/Free task, as animals were permitted to interact with the nose-poke hole and lever during the ITI before orienting and approaching the magazine. Only the second approach and subsequent attempts were included in the analysis. Dopamine traces were then analyzed for these specific moments.

#### Characterization of No-go behaviors

Behavioral strategies employed by rats during No-go trials were manually scored by an experimenter to systematically assess four main features: vigor (scored 0 to 2), and binary indicators for turning, digging, and biting. To characterize vigor, this was based on a scale of 0 to 2: *0* indicating that the animals were not moving, *1* indicating some energy in their movement, and *2* indicating vigorous movement. Turning was defined as the rat rotating the axis of its body while maintaining their snouts in the nose-poke port. Digging was defined as the repeated placement of their paw(s) into the nose-poke hole. Biting was defined by biting the wall of the nose-poke port.

### 6. Statistical Analysis

Data were processed using custom scripts written in MATLAB and Python, and only correct trials were included. Behavioral data were analyzed using GraphPad Prism (v 10.3.1). Due to violations of normality assumptions, a non-parametric Wilcoxon matched-pairs signed rank test was used for pairwise comparisons (trial type) within each task. Kruskal-Wallis tests were used to examine differences across three trial types, and statistical significance was followed-up with *post hoc* pairwise comparisons using Dunn’s test. All tests were two-tailed and statistical significance was set at *p* < 0.05.

#### Dopamine time-series analysis

Dopamine traces within each recording session were directly compared against each other by bootstrapping the within-subject difference trace (e.g., mean Go trace - mean No-go trace within-subject; Jean-Richard-dit-Bressel et al., 2022). Bootstrapped means were obtained by randomly resampling from subject mean dopamine traces with replacement (1000 iterations). CI limits were derived from 2.5 and 97.5 percentiles of bootstrap distribution, expanded by a factor of 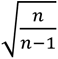 (Jean-Richard-dit-Bressel et al., 2020). Significant transients were identified as a period that CI limits did not contain 0 for at least 1 sec (Jean-Richard-dit-Bressel et al., 2020). To determine the timing of maximum dopamine release for realigned traces, we searched for the maximum dopamine concentration −2 and +2 s around magazine arrival.

#### Clustering No-go behaviors and regrouping animals

Prior to clustering, manually scored trial features were normalized. Averages of characterized No-go scores from each animal across all sessions were then clustered into three groups using agglomerative hierarchical clustering with Ward’s linkage method and Euclidean distance as the metric. A decision-tree classifier was implemented in MATLAB to classify labelled trials into their respective groups, and digging and biting were key features identified for group classification (Supplementary Figure S4). For Go/No-go tasks, average dopamine was calculated within a 2s window following action-cue onset; and for Go/No-go/Free tasks, within 3s after action-cue onset. Similarly, to determine maximum dopamine release following action-cue offset, values were determined within 2s for *Go/No-go* tasks and 3s for Go/No-go/Free tasks. Individual trial data were statistically compared using Kruskal-Wallis tests and statistical significance followed-up with *post hoc* Dunn’s test.

An additional analysis was performed in which animals were regrouped based on their predominant action suppression style within the task (i.e., *Biting*, *Digging* or *Calm*). Regrouped animals were used to compare average and maximum dopamine responses across trial types and in response to unpredicted pellets. Due to limited sample sizes (≤ 2) within certain subgroups, the resulting data were qualified.

**Table 1.** Number of subjects per task variant with FSCV recordings and number of trials in each No-go behavioral classification. Statistical analyses were performed between classified No-go *trials* due to limited sample sizes per group.

|  |  | Total (Biting, Digging, Calm) |  |
|---|---|---|---|
| **Task variant** | **Animals with FSCV** | **Trials** | **Animals with video** |
| Self-initiated Go/No-go | **9** | **229** (61, 89, 79) | 8 (2, 3, 3) |
| Cue-initiated Go/No-go | **11** | **225** (57, 64, 104) | 11 (2, 4, 5) |
| Self-initiated Go/No-go/Free | **5** | **117** (41, 15, 61) | 5 (2, 1, 2) |
| Cue-initiated Go/No-go/Free | **15** | **284** (57, 127, 100) | 14 (3, 6, 5) |

## Acknowledgements

We would like to thank members of the Willuhn laboratory for productive discussions. We thank R. Hamelink and A.S.C. França, for providing technical assistance; M.R. Kandroodi for advice on the hierarchical analysis; L. Fellinger, F. Veen, and T. Arbab for input on the manuscript; H.E.M. den Ouden for constructive feedback. This research was supported by a VICI grant from the Netherlands Organization for Scientific Research VI.C.222.052 awarded to I.W. Funding from the European Union’s Horizon 2020 research and innovation programme under the Marie Skłodowska-Curie grant agreement No 101066959 was awarded to E.Z.P.

## Author contributions

E.Z.P and I.W designed the experiments; E.Z.P, N.B., G.H., L.M., and P.G. performed experiments; N.B., and G.Z., processed video data, E.Z.P. preprocessed and analyzed all data; I.W., L.M., G.H., and P.G assisted with data analysis; E.Z.P. and I.W. wrote and edited the manuscript.

