## Supplementary material for "Ventromedial striatal dopamine dynamically integrates motivated action and reward proximity"

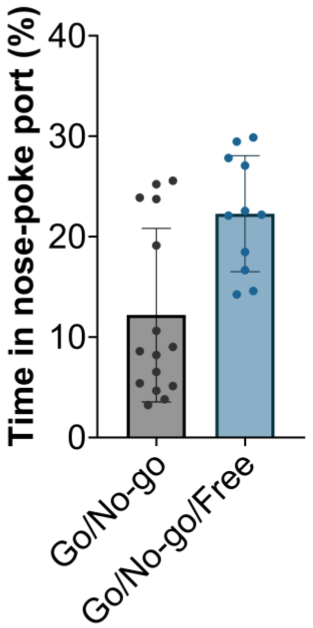


**Supplementary Figure S1. Rats did not continuously occupy the nose-poke port during self-initiated intertrial intervals (ITIs).** Percentage of time spent in the nose-poke port between trials of *self-initiated* Go/No-go and Go/No-go/Free task variants. Animals were permitted to enter the nose-poke port during the ITI but entry could not trigger action-cue presentation. The ITI for the *self-initiated* Go/No-go task was fixed at 5s, whereas for Go/No-go/Free task, it was varying between 15-25s.

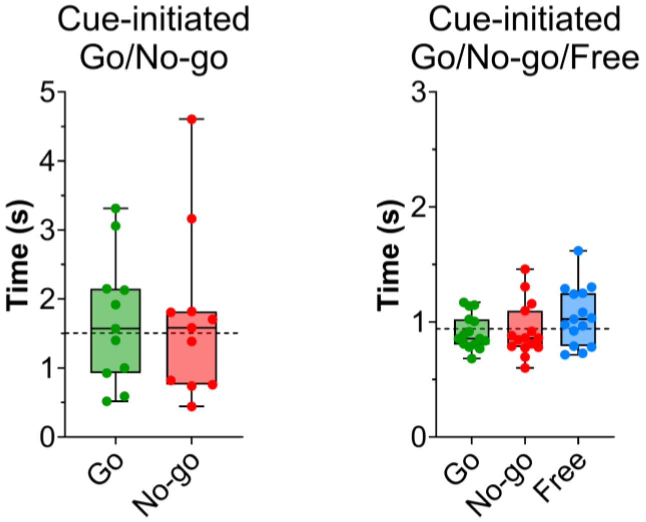


**Supplementary Figure S2. Latency to initiate trials in *Cue-initiated* task versions.** The latency to initiate a trial **following nose-poke light illumination** was similar across trial types within task variants. *Cue-initiated* Go/No-go: W = 18, p = 0.47; *Cue-initiated* Go/No-go/Free: H(2) = 0.93, p = 0.63. Horizontal dotted lines depict the median response latency to start trials after the cue turned on.

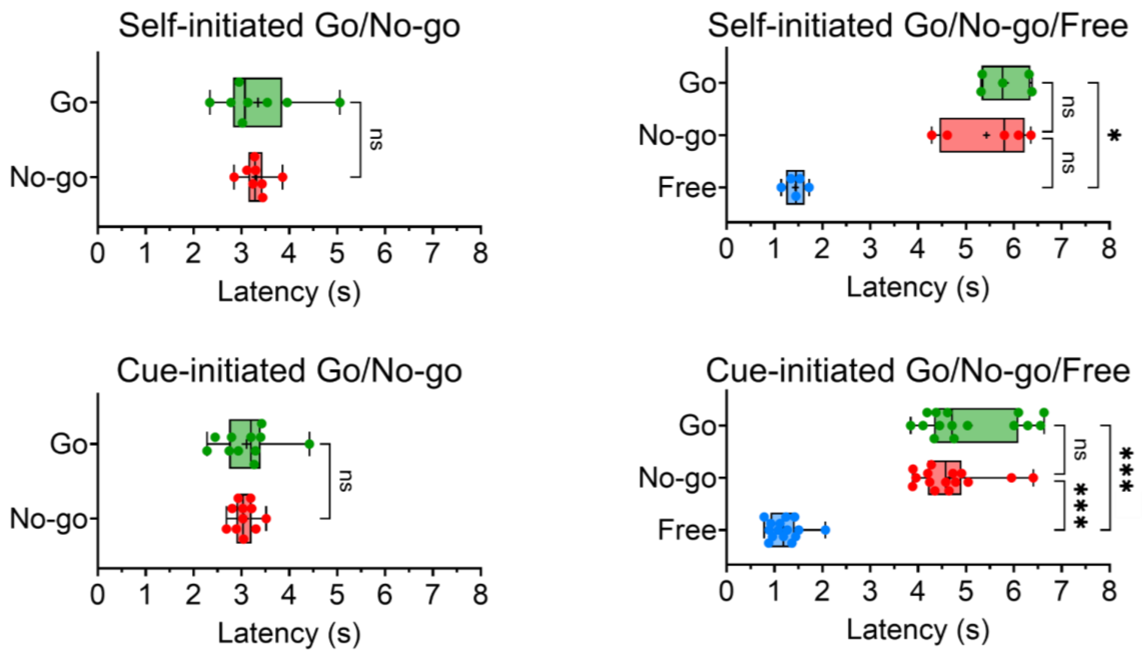


**Supplementary Figure S3. Latency to arrive at the reward magazine was not significantly different between Go and No-go trials.** After satisfying response requirements in Go and No-go trials, rats arrived at the reward magazine at similar time points. During Free trials, animals arrived at the magazine within 2s of hearing the action-cue tone. Action-cue onset is at 0s. Significance for *post hoc* Dunn's tests: * p < 0.05, ** p < 0.01, *** p < 0.001.

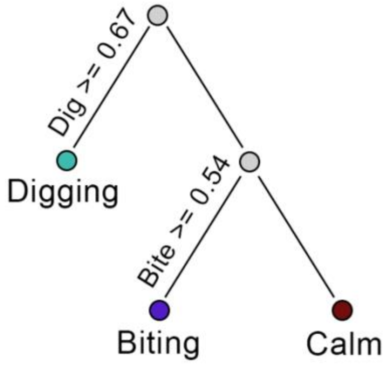


**Supplementary Figure S4. Decision-tree classification of No-go trials.** Individual trials (*n* = 855) were categorised into one of three groups.

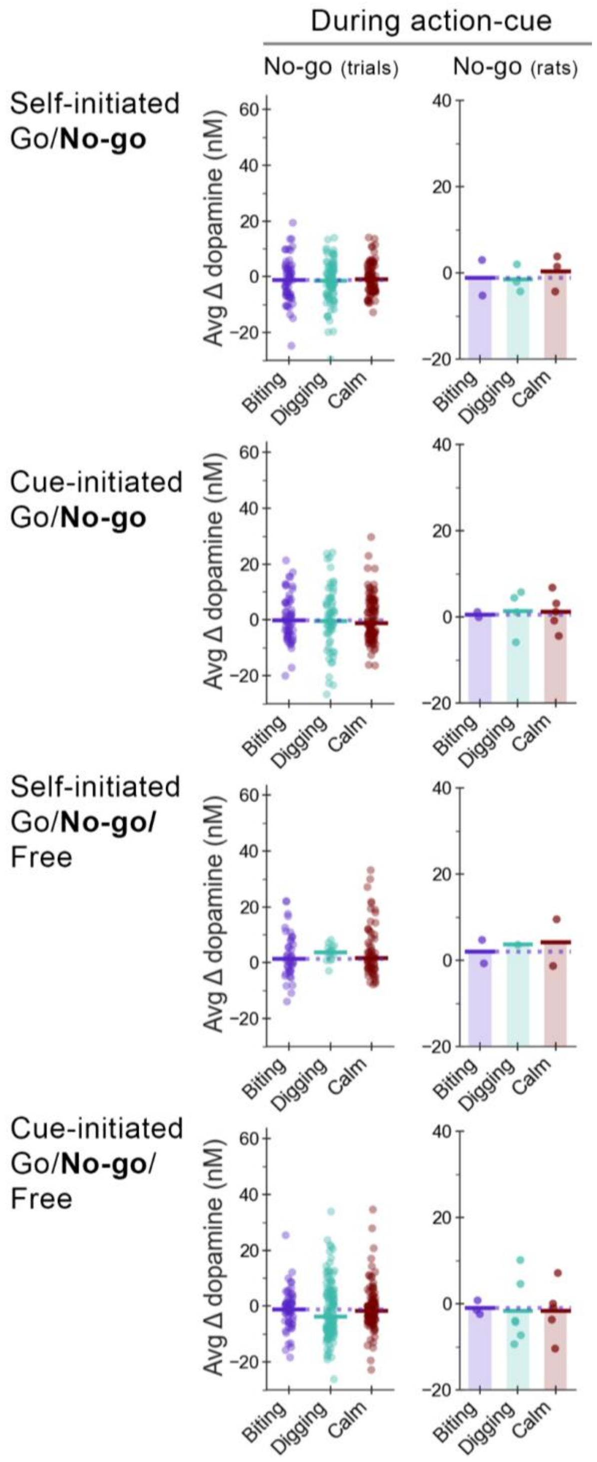


**Supplementary Figure S5. Classified No-go behaviors reveal no changes in dopamine during the action-cue epoch.** *Column 1*) Dopamine concentration during action-cue presentation did not differ significantly between no-go trial classifications (all $p > 0.05$; Kruskal-Wallis test statistics in Statistics table). Solid horizontal lines depict the median. *Columns 2-4)* Rats were grouped according to their predominant No-go strategy (see Methods; horizontal lines depict the mean). *Column 2)* Differences in average dopamine in No-go animals did not qualitatively differ between groups.

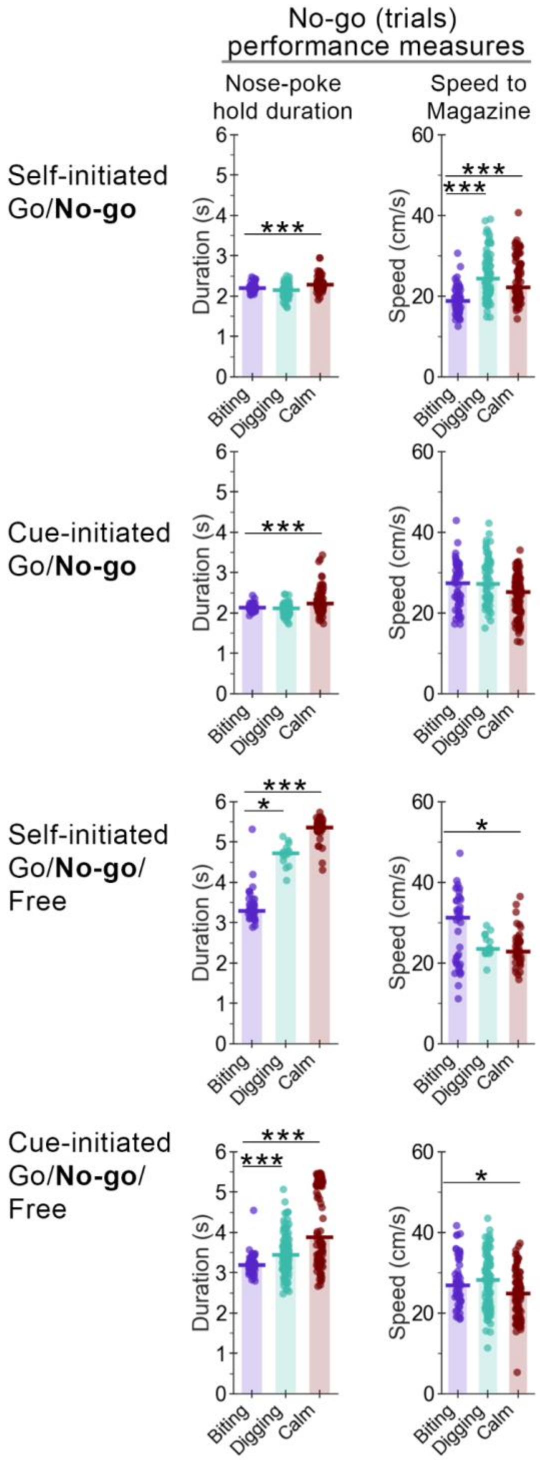


**Supplementary Figure S6. *Biting* trials were characterized by earlier nose-poke exits but inconsistent magazine approach speeds, ruling out a generalized vigor explanation for reward-approach dopamine changes.** *Column 1)* Latency to exit the nose-poke port from action-cue onset. Across Go/No-go/Free task variants, *Biting* trials consistently exhibited the shortest nose-poke hold times. *Column 2)* Speed to approach the reward magazine after nose-poke exit. While Kruskal-Wallis tests were statistically significant for all comparisons ($p < 0.05$; see supplementary statistics table), magazine approach speed showed no consistent relationship with *Biting* trials relative to other classifications. Together, the results rule out 'vigor' during magazine approach as the sole driver of reward-approach dopamine changes. *Post hoc* Dunn's test significance: * $p < 0.05$, ** $p < 0.01$, *** $p < 0.001$.

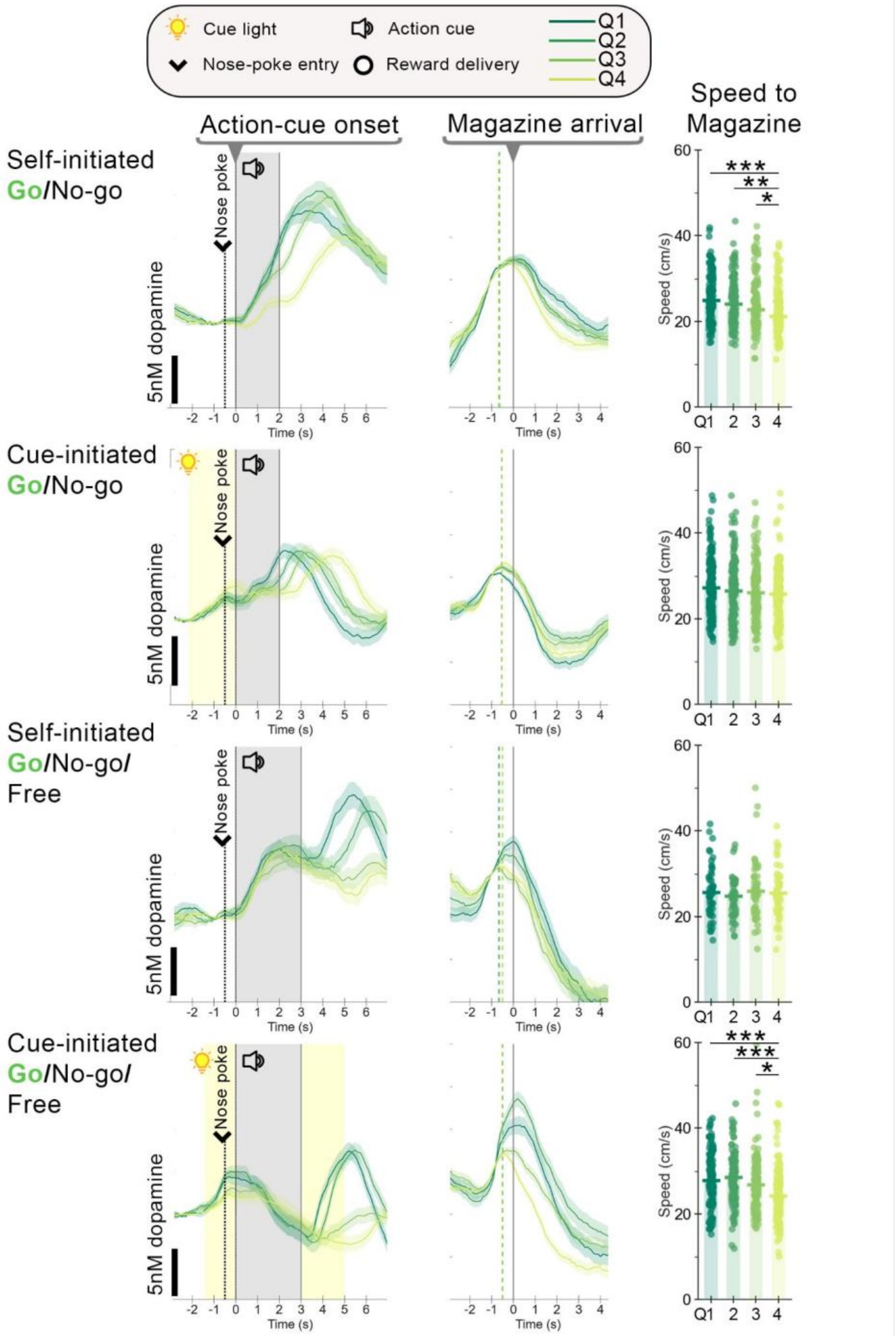


**Supplementary Figure S7. VMS dopamine during Go trials tracks magazine arrival after the final lever-press and is independent of vigor.** For each task variant, trials were divided into quartiles based on last lever-press latencies (Q1 = fastest 25%; Q4 = slowest 25%). *Column 1)* When aligned to action-cue onset, the timing of maximum dopamine during reward approach differed between quartiles. *Column 2)* When realigned to magazine arrival, maximum dopamine was aligned to magazine arrival rather than lever press, as evidenced by consistent timing of maximum dopamine across quartiles. *Column 3)* Speed to approach the reward magazine after the last lever press. Kruskal-Wallis tests were statistically significant for Self-initiated Go/No-go and Cue-initiated Go/No-go/Free ($p < 0.001$; see supplementary statistics table). Comparisons of magazine approach speed within each task variant showed no consistent relationship between quartiles, ruling out post-action-cue 'vigor' as a main driver of reward-approach dopamine changes. *Post hoc* Dunn's test significance: * $p < 0.05$, ** $p < 0.01$, *** $p < 0.001$.

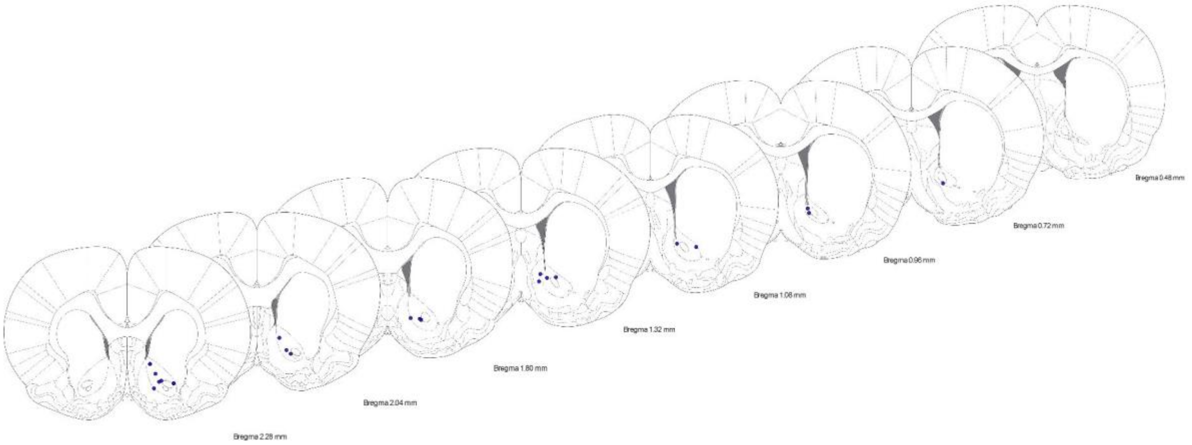


**Supplementary Figure S8. Schematic of all VMS dopamine recording sites.** Coronal sections range from 0.48 to 2.28 mm anterior to Bregma. (Paxinos & Watson 1998).

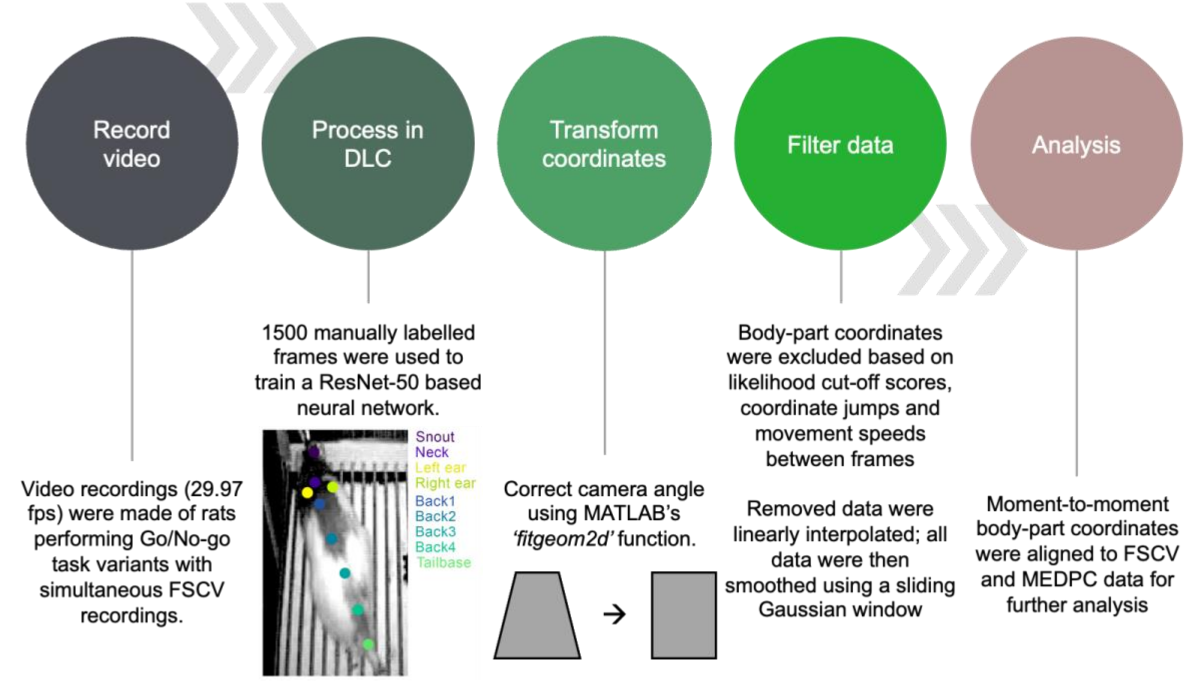


**Supplementary Figure S9. Video analysis workflow.** Details for each step can be found in *Methods Section 5: Video recordings and analysis - DeepLabCut pose estimation*. Analysis details can be found under *DeepLabCut behavior analysis*.

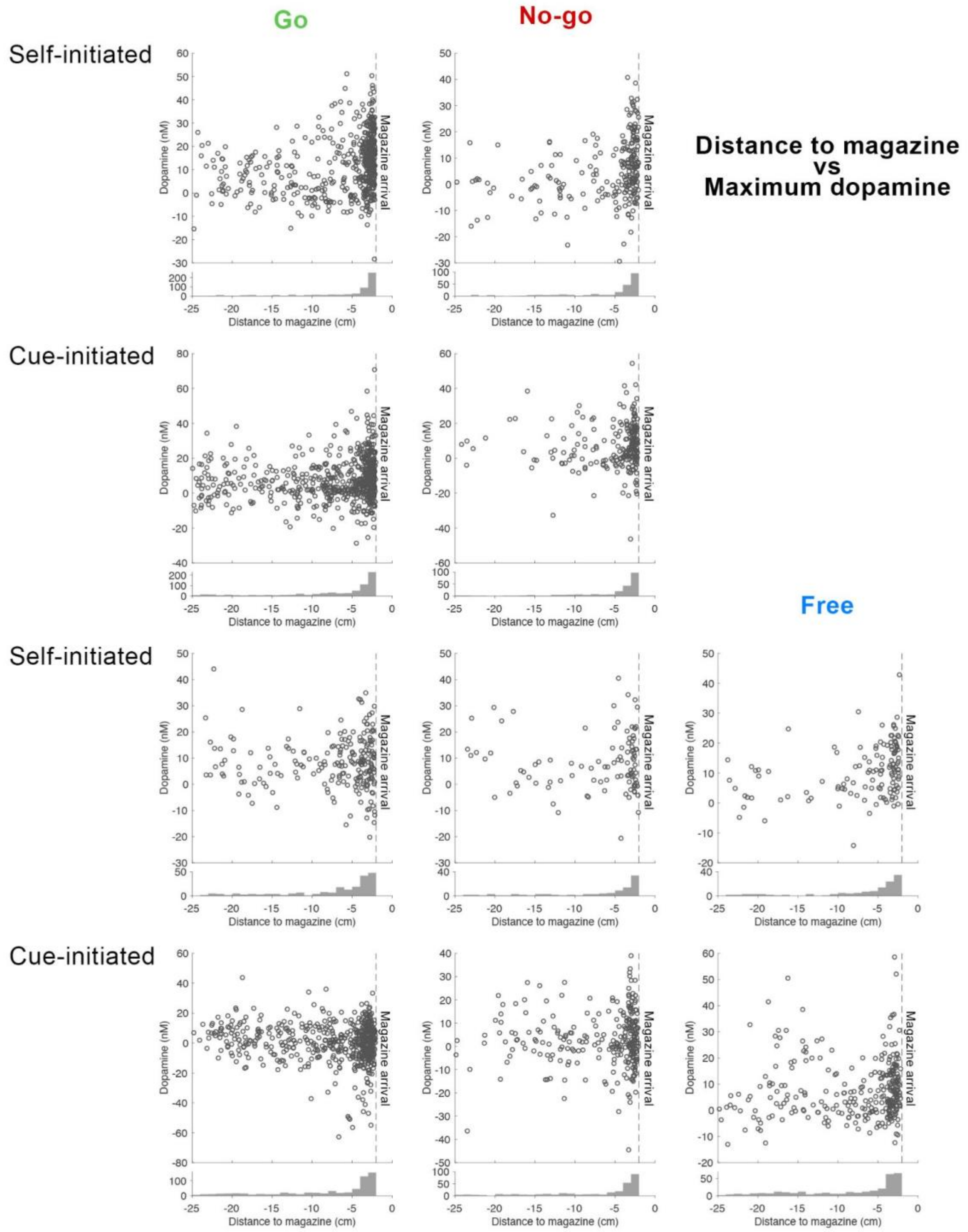


**Supplementary Figure S10. Maximum dopamine concentration per trial as a function of distance to the magazine.** The x-axis represents neck distance from the magazine, progressing from furthest to closest (-2cm). Magazine arrival was defined as the moment neck distance reached <2 cm from the magazine-panel (dotted vertical line). The y-axis shows maximum dopamine concentration recorded within each trial. Individual dots represent the maximum dopamine concentration per trial. Bottom insert (histogram) shows the distribution of trial counts across distance bins. Note the accumulation of max dopamine values as animals arrive at the magazine (dotted line).

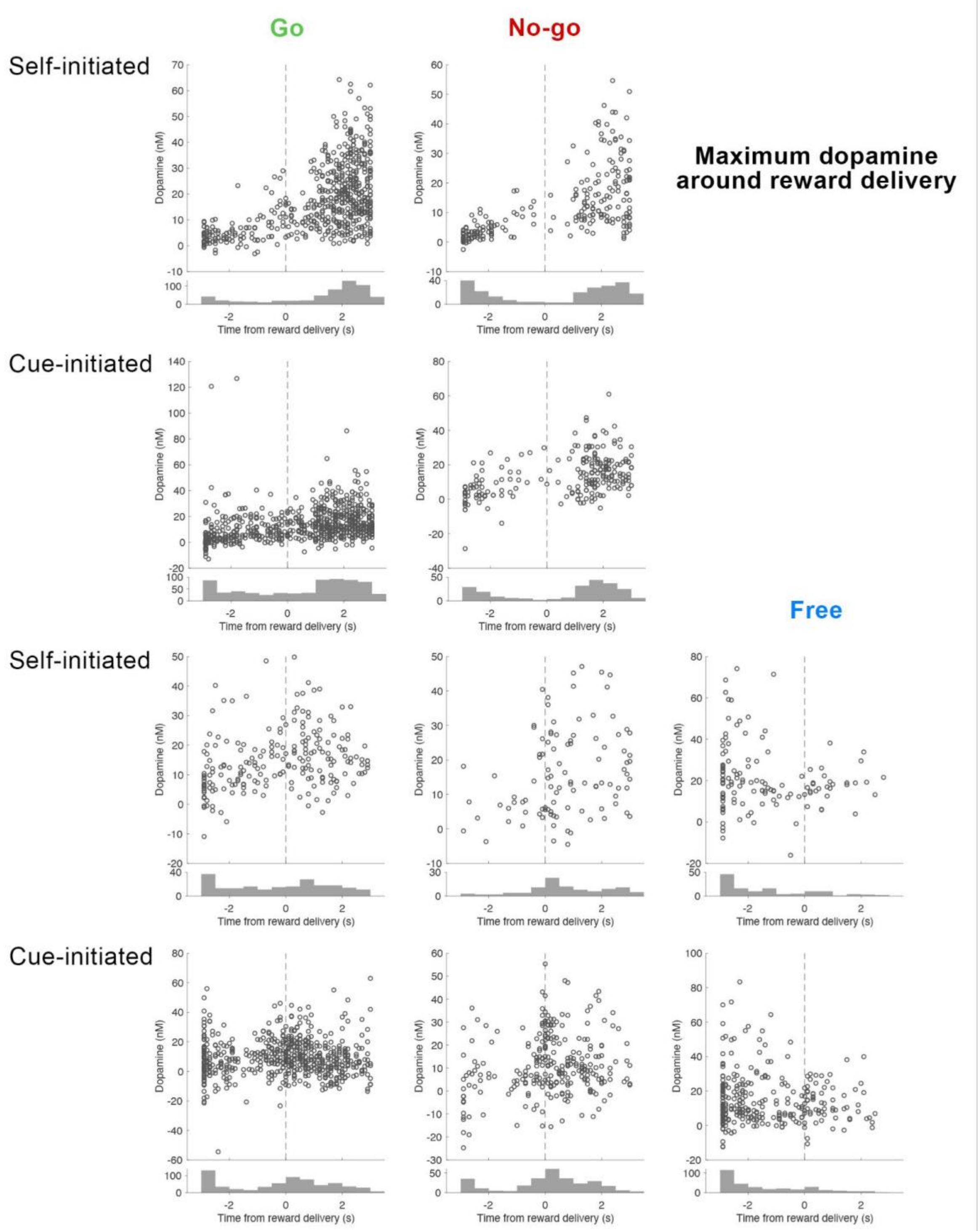


**Supplementary Figure S11. Maximum dopamine concentration around reward delivery (±3s).** The x-axis represents time (s), centered on reward delivery (time = 0 s). The y-axis shows maximum dopamine concentration recorded within each trial. Individual dots represent the maximum dopamine concentration per trial. The bottom insert (histogram) shows the distribution of trial counts across time bins. Note the accumulation of peak dopamine values prior to reward delivery in Free trials but not the other trial types.

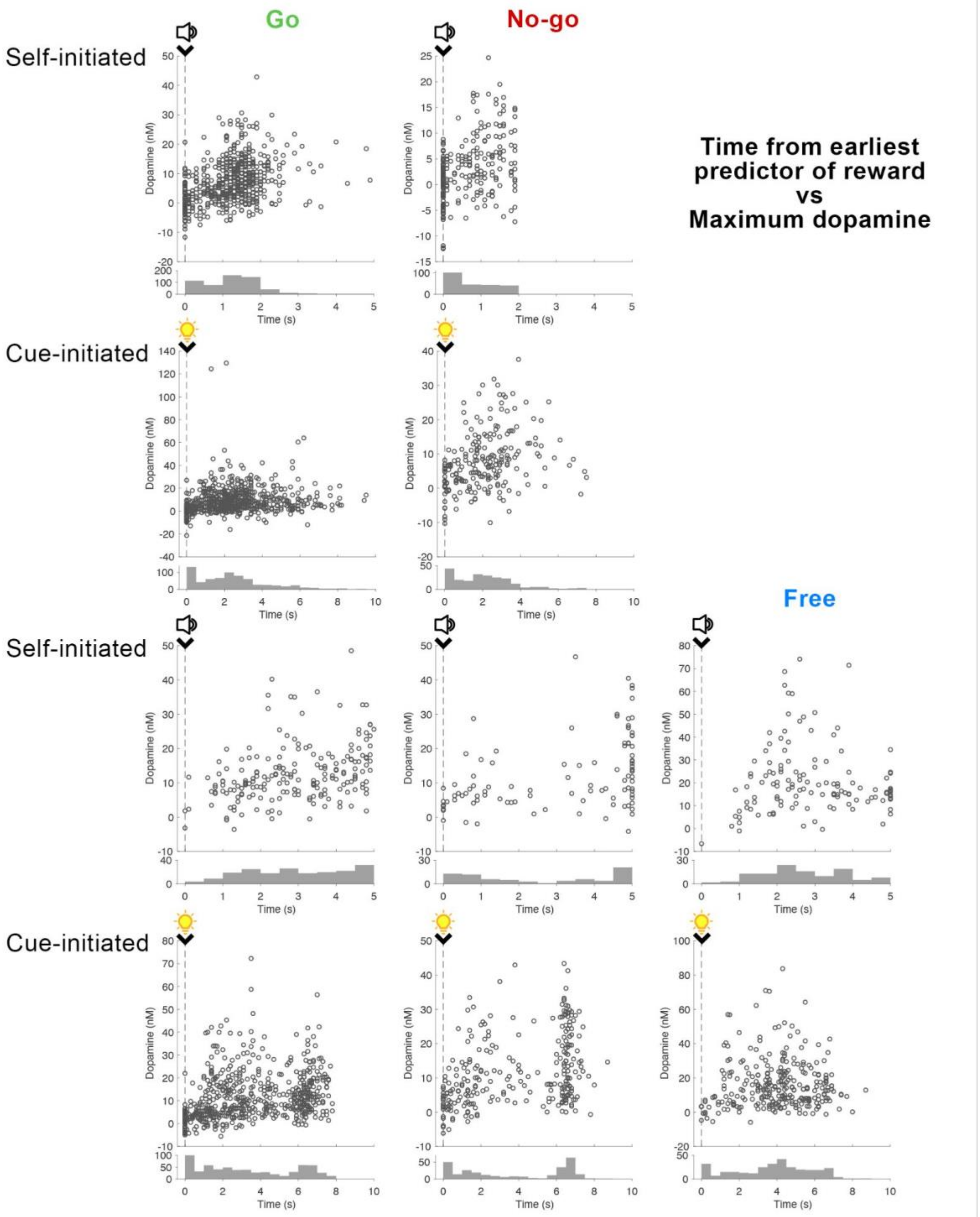


**Supplementary Figure S12 Maximum dopamine concentration from the earliest predictor of reward to reward delivery.** The x-axis represents time (s), with time = 0 s depicting the earliest predictor of reward in each task variant. The y-axis shows maximum dopamine concentration recorded within each trial. Individual dots represent the maximum dopamine value per trial. The bottom insert (histogram) shows the distribution of trial counts across time bins. For *Self-initiated* task variants, time = 0 s corresponds to the onset of the action cue (speaker symbol). For *Cue-initiated* task variants, time = 0 s corresponds to illumination of the nose-poke light (lightbulb symbol). Note that for *Cue-initiated* Go/No-go/Free, 2 Go trials and 3 Free trials fell outside the x-axis range (0–10 s); for Self-initiated Go/No-go/Free, 7 Go trials and 10 Free trials fell outside this range.
